# Contribution of carbon inputs to soil carbon accumulation cannot be neglected

**DOI:** 10.1101/2023.07.17.549330

**Authors:** Xianjin He, Rose Abramoff, Elsa Abs, Katerina Georgiou, Haicheng Zhang, Daniel S. Goll

## Abstract

In a recently published paper (1), the authors report that microbial carbon use efficiency (CUE) is the primary determinant of global soil organic carbon (SOC) storage and that the relative impact of plant carbon inputs on SOC is minor. While soil microbes undoubtedly play an important role in soil organic carbon cycling, we are concerned about the robustness of the approach taken by Tao et al. (1) and highlight potential biases in their analyses that may lead to misleading, model-dependent results.

---

In a recently published paper^1^, the authors report that microbial carbon use efficiency (CUE) is the primary determinant of global soil organic carbon (SOC) storage and that the relative impact of plant carbon inputs on SOC is minor. While soil microbes undoubtedly play an important role in soil organic carbon cycling, we are concerned about the robustness of the approach taken by Tao et al.^1^ and highlight potential biases in their analyses that may lead to misleading, model-dependent results.

An important piece of evidence in support of an empirical relationship between CUE and SOC stems from a meta-analysis based on 132 paired CUE and SOC measurements. Tao and colleagues^1^ applied a linear mixed effect model to this dataset that included CUE, mean annual temperature (MAT), soil depth, and random effects, and explained 55% of the variation in the log-transformed SOC (Fig. 2a and Extended Data Table 1 in Tao et al.^1^). In their linear mixed effect model, carbon inputs to soil were not included despite the authors acknowledging past empirical and theoretical evidence for a major role. To demonstrate that C inputs can also drive SOC variation in their dataset, we extracted net primary production (NPP) from the globally gridded MODIS^2^ for each soil sampling location and used it as a first-order proxy for soil carbon inputs following ref. ^1^. By replacing CUE with NPP in the authors’ linear mixed effect model, we explained a larger proportion of the variation in SOC – namely, 71% with NPP compared to 55% with CUE (Table 1 in the present manuscript). This finding suggests that the empirical results of Tao et al.^1^ may not be robust to the inclusion of other variables, and raises questions about the importance of CUE in explaining SOC variations.

**Table 1.**
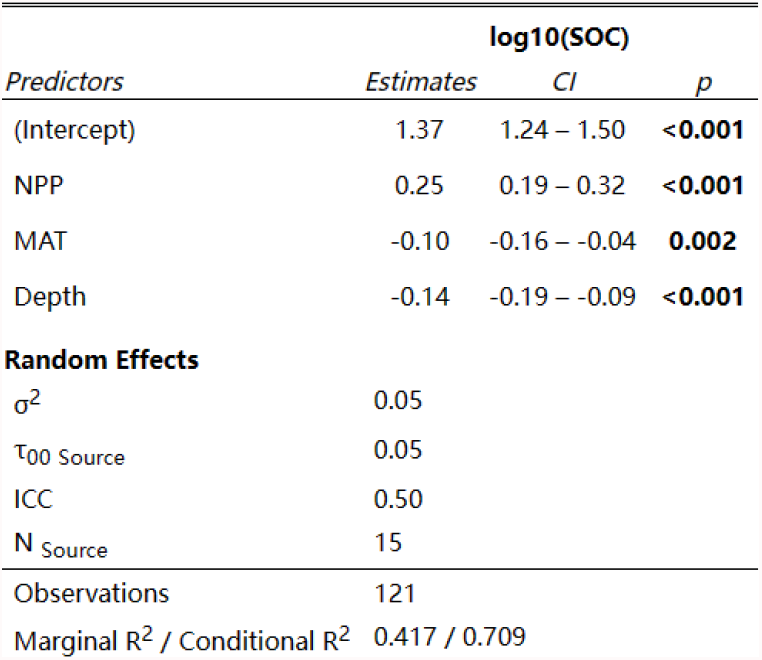
Net primary productivity may explain more variation in soil organic carbon storage than carbon-use efficiency. We performed the same mixed-model regression analyses as in Tao et al., but also explored the importance of net primary production (NPP; g C m^-2^ yr^-1^) as a first-order proxy for carbon inputs to the soil. In both the present study and in Tao et al. (2023), the linear mixed effect model also includes mean annual temperature (MAT; °Celcius) and soil depth (cm), and the study sources were added as the random effects. To ensure the comparability of coefficients across all three explanatory variables (i.e., NPP, MAT, and depth) in the results, we applied standardization using the Z-score method, which maintains the explanatory power of the model.

| *Predictors* | **log10(SOC)** |  |  |
| --- | --- | --- | --- |
|  | *Estimates* | *CI* | *p* |
| (Intercept) | 1.37 | 1.24 – 1.50 | **<0.001** |
| NPP | 0.25 | 0.19 – 0.32 | **<0.001** |
| MAT | -0.10 | -0.16 – -0.04 | **0.002** |
| Depth | -0.14 | -0.19 – -0.09 | **<0.001** |
| **Random Effects** |  |  |  |
| $\sigma^2$ | 0.05 | | |
| $\tau_{00}$ Source | 0.05 | | |
| ICC | 0.50 |  |  |
| N<sub>Source</sub> | 15 |  |  |
| Observations | 121 |  |  |
| Marginal R<sup>2</sup> / Conditional R<sup>2</sup> | 0.417 / 0.709 |  |  |

Tao et al. (2023) further present results from a parameter sensitivity analysis of a process-oriented model which showcase a causal and dominant relationship between CUE and SOC (Fig. 4 in Tao et al., 2023). In order to address uncertainties in model structure and parameters that hamper robust model predictions, the authors used a comprehensive model-data assimilation approach to calibrate a selection of 23 parameters of a SOC model based on a global dataset of soil organic carbon measurements. The calibrated SOC model was then used to quantify the sensitivity of SOC predictions to a selection of potential drivers of SOC, i.e., by varying their values around the optimal or prescribed values one-by-one. We argue that the omission of C inputs and a microbial parameter shown to critically affect the sensitivity of SOC to changes in C inputs in microbial explicit SOC models in the set of optimized parameters raises doubts about the robustness of the findings of the sensitivity analysis.

First, Tao et al. (2023) assumed a SOC model structure which may inherently predispose their analyses to suggest a low importance of C inputs on steady-state SOC. In particular, the chosen model represents the rate of microbial turnover as a linear function of microbial biomass (i.e., ‘density-independent’ with exponent β=1) as opposed to a potential super-linear function (i.e., ‘density-dependent’ with β>1) as suggested in past studies^3–5^. Without this density-dependent microbial turnover, a given change in C inputs may result in a proportional change in the microbial biomass pool and a consequent insensitivity of the SOC pool. This type of model is inconsistent with multiple empirical and theoretical results which show that steady-state SOC pools are sensitive to changes in C inputs, and that this can be better simulated using SOC models with density-dependent microbial turnover^3^. Figure 1 in the present manuscript shows that a switch from density-independent (β=1) to density-dependent (β>1) microbial turnover greatly increases the impact of C input to SOC in the MIcrobial-MIneral Carbon Stabilization (MIMICS) model^6^ (Fig. 1a-c) and in the Millennial model^4^ (Fig.1d-f).

**Fig. 1.**
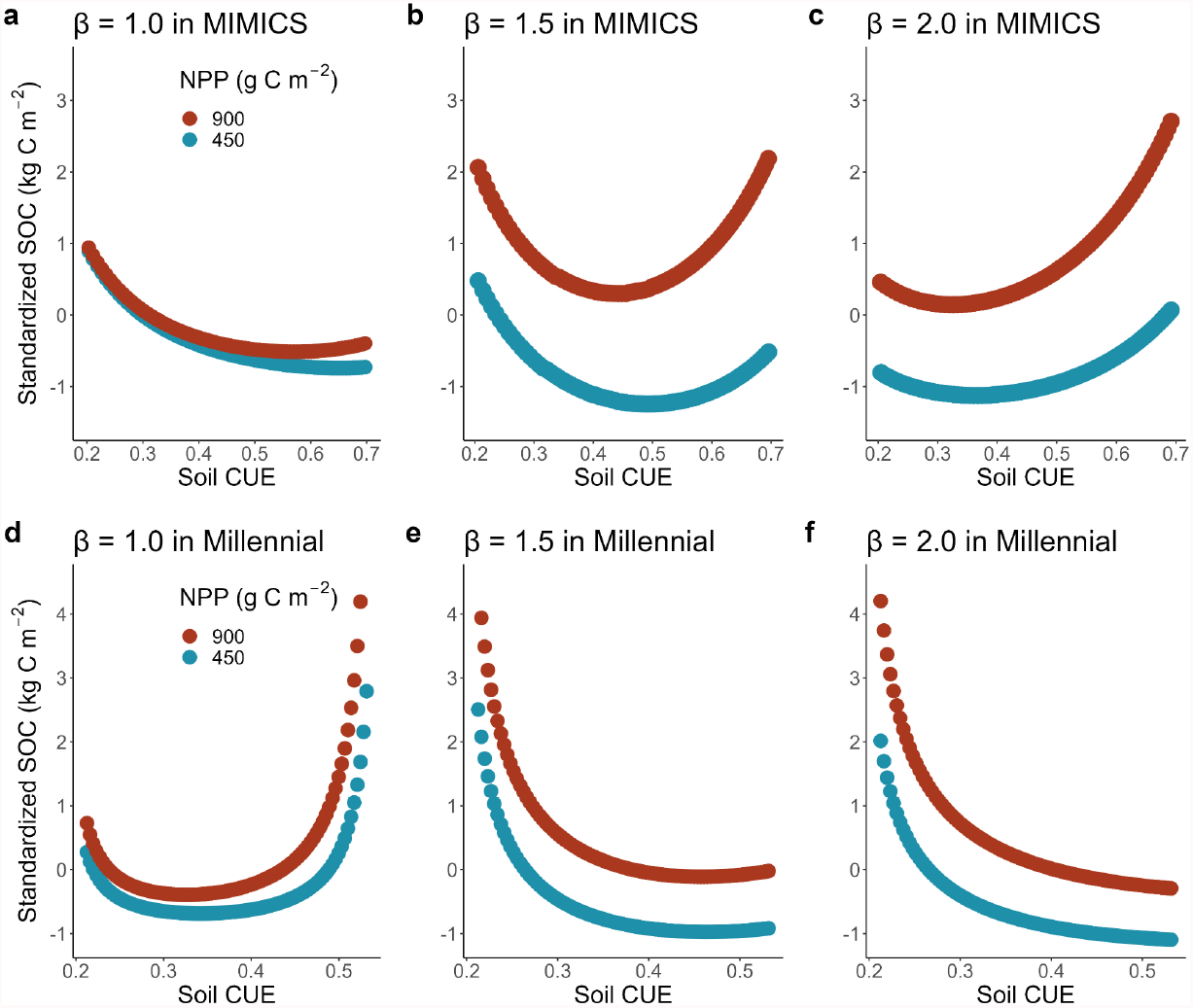
Sensitivity of the CUE-SOC relationship to the inclusion of density-dependent microbial turnover in process-based soil models. Predicted soil organic carbon (SOC) stocks at steady-state from the MIMICS (a-c) and MILLENNIAL (d-f) microbial-explicit SOC models using a range of density-dependent microbial turnover exponent (β) values, net primary productivity (NPP), and microbial carbon use efficiency (CUE). Simulations for a mean annual temperature of 20°Celsius, soil clay content of 20%, and litter lignin to nitrogen ratio of 10. The SOC values in each plot were standardized using the Z-score method to ensure comparability.

While Tao et al. explored the potential need for a sub-linear exponent on the rate of enzyme production –i.e., enzyme production ∼ (microbial biomass)^β^ where 0< β< 1– in their SOC model, this modification is functionally and theoretically distinct from the density-dependent microbial turnover with β>1 proposed in earlier work^3^. We conducted a sensitivity analysis^7^ to determine whether SOC behaved the same if a β exponent was assigned to enzyme production (0< β< 1 as in^1^) versus microbial turnover (1< β< 2 as in^3^). We found that the sensitivity of SOC to a variation of +/-10% of CUE is equal to 1.3 when β=1 irrespectively of where it is assigned, but is much less when β is not equal to 1: 0.48 and 0.73 for a 50% change in β on enzyme production and turnover, respectively. On the other hand, the sensitivity of SOC to a variation of +/-10% of C input is equal to 0 when β=1, 0.52 when β is modified by 50% on enzyme production, and 0.34 when β is modified by 50% on turnover. This indicates that the results of Tao et al. are very contingent on the assumed model structure. If β associated with turnover is not found with observations to be mostly 1 (as for enzyme production), then a lower sensitivity of SOC to CUE and a greater sensitivity of SOC to C input may have been observed. Besides, the exploration of β by Tao et al. are only in the reply to the reviewers and it is not sufficiently described how the results were obtained.

Second, Tao et al. approximated C inputs to the soil using net primary production (NPP) from predictions of a land surface model. NPP is a notoriously uncertain carbon flux and it is not clear to what extent NPP from land surface models actually reflects C inputs to soil and its spatial variations^8^. The use of the interannual variation in NPP from a single land surface model to characterize uncertainty in C inputs, as done in the optimization in this study, falls arguably short to characterize the true uncertainty. Its implications for the outcome of the study remains elusive, representing a source of uncertainty. The inclusion of C input^9^ as a parameter for optimisation at site rather than the inclusion of NPP as environmental driver for the global extrapolation^1^ of site-specific optimized parameters could be a way forward.

In summary, we highlight several statistical and process-based model assumptions that may have biased the overarching conclusion that CUE is the dominant control on spatial variation of SOC. We argue that changes in soil microbial CUE itself are influenced by environmental factors, including carbon inputs as well as the quality of litter ^10,11^. Tao et al.’s findings contradict numerous empirical studies that report that changes in plant inputs significantly alter soil carbon (e.g., in ^12–14^). We believe that further examination of statistical and process-based model structures is needed to demonstrate the robustness of the conclusions presented.

## Acknowledgement

DSG and RA acknowledge the generosity of Eric and Wendy Schmidt by recommendation of the Schmidt Futures program, DSG and XH acknowledge support from EJP Soil ICONICA project. Funding to EA was provided by the European Union’s Horizon 2020 research and innovation programme under the Marie Sklodowska-Curie grant agreement No 891576.

## Contributions

DSG, XH and EA conceptualized and designed this idea. XH, RA, EA, KG, HZ, and DSG discussed the results and contributed to the text.

